# Rift Valley fever in northern Senegal : a modelling approach to analyse the processes underlying virus circulation recurrence

**DOI:** 10.1101/2019.12.23.886978

**Authors:** B. Durand, M. Lo Modou, A. Tran, A. Ba, F. Sow, J. Belkhiria, A.G. Fall, B. Bitèye, V. Grosbois, V. Chevalier

## Abstract

Rift Valley fever (RVF) is endemic in northern Senegal, a Sahelian area characterized by a temporary pond network that drive both RVF mosquito population dynamics and nomadic herd movements. To investigate the mechanisms that explain RVF recurrent circulation, we modelled a realistic epidemiological system at the pond level integrating vector population dynamics, resident and nomadic ruminant herd population dynamics, and nomadic herd movements recorded in Younoufere area [1]. To calibrate the model, serological surveys were performed in 2015-2016 on both resident and nomadic herds in the same area. Mosquito population dynamics were obtained from a published model trained in the same region [2]. Model comparison techniques were used to compare five different scenarios of virus introduction by nomadic herds associated or not with vertical transmission in *Aedes vexans*. Our serological results confirmed a long lasting RVF endemicity in resident herds (IgG seroprevalence rate of 15.3%, n=222), and provided the first estimation of RVF IgG seroprevalence in nomadic herds in West Africa (12.4%, n=660). Multivariate analysis of serological data suggested an amplification of the transmission cycle during the rainy season with a peak of circulation at the end of that season. The best scenario of virus introduction combined yearly introductions of RVFV from 2008 to 2015 (the study period) by nomadic herds, with a proportion of viraemic individuals predicted to be larger in animals arriving during the 2^nd^ half of the rainy season (3.4%). This result is coherent with the IgM prevalence rate (4%) found in nomadic herds sampled during the 2^nd^ half of the rainy season. Although the existence of a vertical transmission mechanism in *Aedes* cannot be ruled out, our model demonstrates that nomadic movements are sufficient to account for this endemic circulation in northern Senegal.

**Author summary:** Rift Valley fever (RVF) is one of the most important vector borne disease in Africa, seriously affecting the health of domestic ruminants and humans and leading to severe economic consequences. This disease is endemic in northern Senegal, a Sahelian area characterized by a temporary pond network that drive both RVF mosquito population dynamics and nomadic herd movements. Two non-exclusive mechanisms may support this endemicity: recurrent introductions of the virus by nomadic animals, and vertical transmission of the virus (i.e. from infected female mosquito to eggs) in local *Aedes* populations. The authors followed up during 1 year resident and nomadic herds. They used the data thus obtained to model a realistic epidemiological system at the pond level integrating vector population dynamics, resident and nomadic ruminant herd population dynamics. They found that the best scenario explaining RVF remanence combined yearly introductions of RVFV by nomadic herds, with a proportion of viraemic predicted to be larger in animals arriving during the 2^nd^ half of the rainy season, which is consistent with an amplification of virus circulation in the area during the rainy season. Although the existence of a vertical transmission mechanism in *Aedes* cannot be ruled out, their results demonstrates that nomadic movements are sufficient to account for this endemic circulation in northern Senegal.

## Introduction

Vector borne pathogen circulation results from complex interactions between hosts, vectors and pathogens. These interactions are modulated by intrinsic factors such as genetic, but also by extrinsic factors such as irrigation, rainfall, human population density, animal transhumance and trade, or social practices [3, 4]. Understanding this complexity is a crucial to control vector borne diseases and mitigate their impacts.

Rift Valley fever (RVF), is one of the most important vector borne disease in Africa, seriously affecting the health of domestic ruminants and humans and leading to severe economic consequences [5, 6]. RVF is an acute, viral disease, caused by a Phlebovirus (Bunyaviridae family) [7]. Infection leads to abortions in pregnant animals and high mortalities in new born sheep and goats. Humans get infected after being in contact with infectious tissues -blood, body fluids or abortion products, or through the bite of an infectious mosquito. Most human cases are moderate, but rare severe complications such as hepatitis or encephalitis could occur and lead to death. Transmission between ruminants occurs via mosquito bites, mainly from *Aedes, Culex* and *Mansonia* genera [8, 9], and probably through direct contact with infectious products [10, 11].

Since its first recognition in 1931 in Kenya, RVF outbreaks have been regularly reported in sub-Saharan Africa [12–18], Indian Ocean [19, 20] and Arabic Peninsula [21]. The most recent outbreaks occurred in Mauritania (2015), Niger (2016), Uganda (2016) and Mayotte (2018-2019) [12, 22–24].

In Senegal, West Africa, RVF is endemic and has been repeatedly reported among humans, livestock, and mosquitoes, especially in the Ferlo (northern Senegal) [25–29]. The Ferlo is a typical sahelian area, with a semi–arid climate, annual rainfall ranging from 300 to 500 mm and a rainy season lasting from July to October/November. This area is sprinkled by a complex and dense network of temporary ponds located within the fossil Ferlo river bed. These ponds fill up during the rainy season and dry out during the rest of the year. During the rainy season, pond water levels show daily fluctuations, increasing with rainfall and decreasing with infiltration (favoured by sandy-loam soils), high evapotranspiration and water consumption by livestock and humans. This water availability combined with high quality grass, attract a massive flow of nomadic herds coming from the whole country, and sometimes from neighbouring countries [1]. Nomadic herds use to settle around these temporary ponds for a couple of weeks or sometimes several months, depending on grass and water availability [1]. In the middle of dry seasons, ponds are totally dried, and only large wells can be used for ruminants.

Temporary ponds are also important breeding sites for RVFV mosquito vectors, i.e. *Culex poicilipes* and *Aedes vexans* [26, 30]. Their respective population dynamics are strongly linked to the pond water surface dynamics, thus to the rainfall pattern [31, 32]. During the first part of the rainy season, sparse rainfall events fill ponds that dry few days later: this succession of filling and emptying stages is favourable for *Ae. vexans* vectors [31]. The second half of the rainy season is characterised by frequent and heavy rainfall events: ponds remain flooded for 2-3 months, allowing the *Culex* population to explode. Rainy season, characterised by high ruminants and RVF vector densities, is thus highly favourable for RVFV transmission. However, during the dry period vectors are absent [31], and the mechanisms allowing RVFV recurrence in this region still remain unknown. As demonstrated in other contexts, movements of viraemic ruminant are likely to contribute to RVFV persistence and spread [33–35]. Belkhiria et al. identified three types of migrations in a recent survey performed in the Ferlo: predominant long-distance country level migrations, short distance migrations limited to the Ferlo region with breeders moving from pond to pond, and inter-countries migrations that extend from Mauritania to Gambia through Senegal [1]. Whatever the distance they travel, these herds pass through infected areas and may transport and introduce the virus into the Ferlo. The second, and still main hypothesis in the literature for persistence of RVFV in the environment between epizootics is vertical transmission in mosquito vectors (VT), *i.e.* the transmission of the virus from infected females to mosquito offspring [36]. This VT mechanism has been demonstrated in Kenya for *Aedes mcintoshi*, and a recent survey carried on in Sudan suggested its existence in *Culex quinquefasciatus*. VT may be an alternative, and non-exclusive, way for RVFV to survive the dry season, and persist in the area despite unfavourable conditions. Although VT has never been demonstrated for *Ae. vexans* in northern Senegal, and although a recent entomological survey showed very low densities of *Ae. mcintoshi* in this area [37, 38], the existence of this mechanism cannot be ruled out. Another mechanism could imply the survival of RVFV in hibernating *Culex* mosquitoes as demonstrated with West Nile virus in the United States [39–41]. However, laboratory and field arguments are lacking and further studies are needed to bring more substantial evidence of these mechanisms. A last putative mechanism implies rodents [42]. However, and despite evidences of association between rodents and RVFV circulation, their potential implication in RVFV maintenance remains controversial [43]. Because of the absence of mosquitoes during the dry season in the study area, it is unlikely that rodents contribute to maintain the virus during this unfavourable season [42].

In the Ferlo, the set composed of a temporary pond, associated vectors, sedentary ruminants living around, and nomadic herds seasonally settling around, constitutes the elementary unit of the RVF epidemiological system. Rainfall variation is the main driver for pond surface fluctuations which affects both the vector population dynamics and nomadic migration patterns. In addition, resident and nomadic herd immunity impact the ability of RVFV to circulate. A better understanding this complex system is needed to improve our comprehension of RVF epidemiology in this region, to quantify the main determinants of RVF transmission and emergence and help establishing better surveillance, prevention and control strategies.

The aim of this work was to model RVFV transmission in this epidemiological system, and to use this model to infer on the respective contribution of nomadic movements and VT in RVFV recurrence. We first carried out in 2016 epidemiological surveys to (i) document the demographic characteristics of both sedentary and nomadic ruminant populations, as well as the duration of nomadic herds stays in our study area, and (ii) estimate the RVFV seroprevalence rate in sedentary small ruminants and nomadic herds transiting through the study area. Then we modelled a realistic epidemiological system at the pond level integrating both *Cx. poicilipes* and *Ae. vexans* population dynamics, resident and nomadic ruminant herd population dynamics, and nomadic herd movements. Vector population dynamics were based on outputs of an entomological model (EM) previously and independently developed, parameterized and validated using mosquito trapping data collected in the same area, during the same period [2]. Model comparison methods allowed analysing the scenarios that could explain the recurrent circulation of the virus in this ecosystem, incorporating VT in *Ae. vexans* or not, direct transmission between hosts or not, and occasional or regular viral introduction through nomadic herd seasonal migrations. The selected scenario was finally used to estimate the transmission parameters based on the observed seroprevalence data, and provide new insights on the role of nomadic herds in RVFV recurrence in the survey area.

## Materials and Methods

### Study area and epidemiological system

The study was conducted in a 86 sq. km area located near Younoufere village (Fig. 1). Located in the Ferlo area, northern Senegal, Younoufere (15.269464° N and 14.463094° W) is 80 km far from Barkedji village, where many epidemiological and/or entomological studies on RVF previously occurred [25, 29, 44–46]. Both areas share the same eco-climatic conditions previously described, i.e. a semi-arid steppe and many temporary ponds filled by seasonal rainfall, and that constitute a perfect habitat for *Cx. poicilipes* and *Ae. vexans* mosquitoes [1, 25, 32, 38].

**Figure 1.**
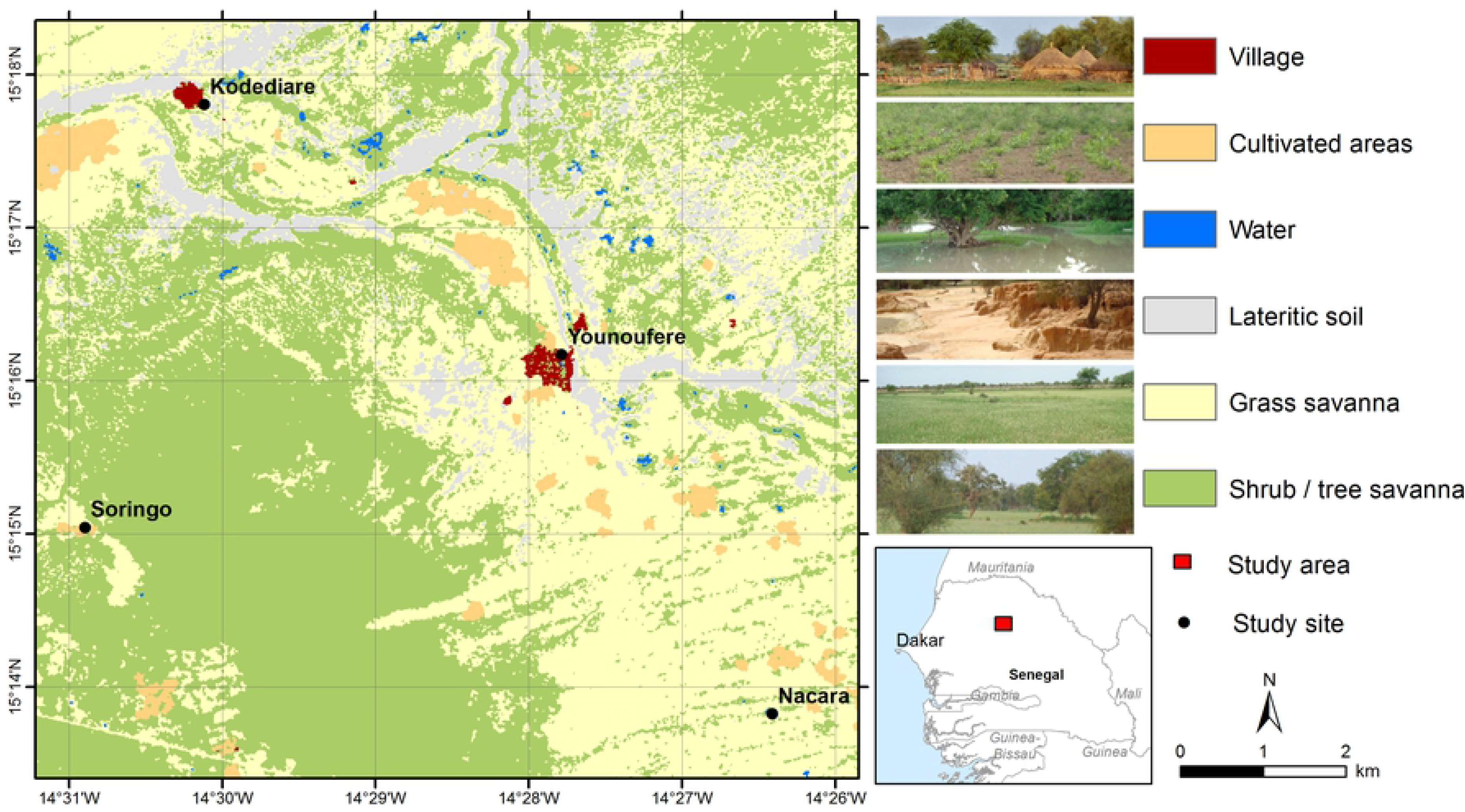
Map of the study area. Map created using ESRI ArcGIS™; land cover map derived from a SPOT-5 image (2014-08-07); administrative boundaries extracted from DIVA-GIS (https://www.diva-gis.org/)

A field systematic census performed during the 2015 rainy season indicated the existence of 25 ponds in the study area, corresponding to an average density of 0.29 pond/sq. km. Younoufere is also a known hub for nomadic herds that may come from the Ferlo area itself, but also from northern or southern regions of Senegal [1]. Upon arrival, nomads use to select a convenient place to settle, never far from a pond, i.e. 100 m to 3 kms [25]. The location of resident herds is also strongly constrained by the availability of livestock watering places. During the rainy season, herds remain located in the neighbourhood of the temporary ponds where sedentary breeders live. During the dry season, sedentary herds have to move every day to distant drilling water place they use as water points.

### Epidemiological surveys

Host population sizes, demographic characteristics/parameters, and serological status of resident and nomadic animals were investigated thanks to field surveys described hereafter.

### Ethical statement

No human experiments were conducted in this study. Meetings were organized with Younoufere villagers and nomadic and resident farmers to explain the goals of the study and the decision to participate was taken at the individual level. Informed consent was given verbally and documented in questionnaires. For cultural reasons, written consent could not be obtained. During this study, we followed the World Animal Health Organisation (OIE) guiding principles on animal welfare included in the OIE terrestrial Code, Chapter 7.8 “Use of Animals in research and education” [47].

### Resident herds population and serological status

There is no existing census of the cattle and small ruminants living in the study area. The average size of sedentary populations was estimated based on the number of animals visiting Younoufere’s well during the dry season. Indeed, temporary ponds are dried out during the dry season and farmers are forced to take their animals to the well every day. Local authorities control access of animals to this watering place, and animal owners that come for watering ruminants are systematically registered. In 2012 dry season–the most recent available registration, an estimate of 6000 cattle and 30,000 small ruminants visited the Younoufere well. Eighty percent of these ruminants, i.e 4800 cattle and 24,000 small ruminants, belonged to resident breeders. We assumed that homes of these resident herders were located near temporary ponds (dried during the dry season) within a 15 km radius of the well. Considering a density of 0.29 pond per sq. km (the average pond density in the study area ‒see above), the expected number of temporary ponds in this 15 km radius area was 205. The average number of sedentary animals living around a given pond was then estimated at 23 cattle (i.e. 4800/205) and 117 small ruminants (i.e. 24000/205).

In May 2016, to assess the serological status of resident herds ling in the study area, a cross sectional serological survey was performed in 6 resident herds living in 3 small villages located in the close vicinity of Younoufere (Kodediare, Nacara and Soringo (Fig. 1)). Animals were randomly chosen among herds whose owner agreed to participate to the study. As a whole, an age-structured sample of 222 small ruminants were blood sampled (no sample from resident cattle could be obtained). Sera were tested by IgG ELISA test using a Kit ID screen©Rift Valley Fever Competition Multi-species from IDvet (Innovative Diagnostics) according to the manufacturer’s instructions. Positive samples were then tested for IgM using the Kit ID screen© Rift Valley Fever IgM capture. Serological analyses were performed in the virological laboratory of ISRA (Dakar, Senegal).

### Nomadic herds population, movements and serological status

The average size of nomadic populations during the dry season was estimated using the data collected at the Younoufere well at 6 cattle and 29 small ruminants in 2012. To estimate the size of nomadic population during the rainy season, we used data from a survey ran between from August 2015 to January 2016 [1]. Briefly, the survey consisted of 6 one week field sessions (4 during the rainy season and 2 in the early dry season) aiming to characterize nomadic movements and their determinants, as well as the RVF serological status of animals when arriving in the study area. The protocol is fully described in [1]. Only one herd was encountered during the last session performed on late January, and this herd could not be included in the sampling frame described below. Session 6 data was then merged with session 5 (Table 1). During each session, the majority of breeders settling in the study area agreed participating to the study and were interviewed.

**Table 1.** Surveys conducted in nomadic breeders of the study area.

| Session | Period | Season | Number of Breeders | Total number of animals |  |  |
| --- | --- | --- | --- | --- | --- | --- |
|  |  |  |  | Cattle | Sheep | Goats |
| 1 | 15/08/2015 to 25/08/2015 | Rainy | 37 | 2591 | 5433 | 1268 |
| 2 | 31/08/2015 to 04/09/2015 | Rainy | 32 | 1801 | 4906 | 1467 |
| 3 | 15/09/2015 to 17/09/2015 | Rainy | 39 | 1589 | 4152 | 1466 |
| 4 | 01/12/2015 to 04/12/2015 | Rainy | 14 | 714 | 2940 | 669 |
| 5 | 09/01/2016 to 11/01/2016 | Early dry | 9 | 640 | 2035 | 370 |
| Average values |  | Rainy | 30.75 | 1673.75 | 4357.75 | 1217.5 |

Information about their arrival and scheduled departure dates were collected [1], which allowed estimating the duration of their stays around a given temporary pond. herders were asked to list all locations where they stayed during the past year as well as all the areas they plan to use during the coming months. For each herd the home location (where the herd usually stayed during the dry season) and the locations visited by the herd during the rainy season were identified. As described in Belkhiria et al.[1], the locations were grouped into three regions: the Northern region (including the Senegal river valley), the Ferlo and the Southern region. This allowed to distinguish two types of nomadic breeders: short-range nomads who stayed yearlong in the Ferlo, moving between temporary ponds during the rainy season, and the long-range nomads, whose home locations were outside the Ferlo (i.e. in Northern or Southern regions) and who visited the Ferlo during the rainy season, moving likewise between temporary ponds. To assess the serological status of nomadic animals entering the study area, 4 herds newly arrived were selected based on owner willingness during each of the 6 sessions. Among the 132 nomadic herders who accepted to answer the above-described movement survey, 22 accepted to participate in a serosurvey and 30 animals (cattle and small ruminants) were randomly sampled from each. Sera were tested for RVF IgG antibodies using the test mentioned above. Positive sera were tested for IgM.

### Statistical analyses

No cattle could be sampled in resident herds and cattle sera were sometimes missing in nomadic herds (Table 2). We thus analyzed the relationship between seropositivity and potential risk factors in small ruminants only, using a logistic mixed model. The outcome was the individual IgG status, and the fixed effects were firstly individual variables: age (years), species (sheep of goat) and sex (male or female). Secondly, resident herds were sampled in one session, in May 2016, whereas nomadic herds were sampled in 5 sessions, between August 2015 and January 2016. The respective effects of the sampling date and of the herd type (short vs long range nomads), could thus not be disentangled. For that reason, we created a composite variable to represent the joint effect of sampling date and herd type. We first defined three groups of sampling dates: (i) the 2015’s rainy season 1^st^ half (i.e. August-September 2015, corresponding to sessions 1-3 in nomadic herds: see Table 1), (ii) the 2015 rainy season’s 2^nd^ half and the early 2016 dry season (i.e. October 2015-January 2016, corresponding to sessions 4-5 in nomadic herds: see Table 1), and (iii) during the 2016 dry season (i.e. May 2016, the sampling period for resident herds). We then defined a composite variable with 5 distinct modalities: short-range (resp. long-range) nomadic herds sampled during the 1^st^ half of the 2015 rainy season (August-September 2015), or during the 2^nd^ half of the 2015 rainy season and the early 2016 dry season (October 2015-January 2016), and resident herd sampled during the 2016 dry season (May 2016). This composite variable was included in the model as a fixed effect. The breeder was treated as a random effect.

**Table 2.** Results of serosurveys conducted in ruminants of resident and nomadic herds in Younoufere area, Ferlo, Senegal, May 2015-January 2016.

| Sampling period | Season | Type of breeder | Sheep | Goats | Cattle |
| --- | --- | --- | --- | --- | --- |
| 15/08/2015 – 25/08/2015 | Rainy (1<sup>st</sup> half) | Short-range nom.<sup>a</sup><br>Long-range nom. | 1/26<sup>b</sup><br>0/40 | 0/13<br>0/15 | 0/21<br>0/5 |
| 31/08/2015 – 04/09/2015 | Rainy (1<sup>st</sup> half) | Short-range nom.<br>Long-range nom. | 3/32<br>4/45 | 5/18<br>1/14 | 3/10<br>0/1 |
| 15/09/2015 – 17/09/2015 | Rainy (1<sup>st</sup> half) | Short-range nom.<br>Long-range nom. | 6/31<br>3/66 | 4/29<br>2/24 |  |
| 01/12/2015 – 04/12/2015 | Rainy (2<sup>nd</sup> half) | Short-range nom.<br>Long-range nom. | 5/26<br>14/69 | 1/13<br>2/9 | 4/21<br>3/12 |
| 09/01/2016 – 11/01/2016 | Early dry | Short-range nom.<br>Long-range nom. | 4/64<br>9/30 | 8/26 |  |
| May 2016 | Dry | Resident | 24/168 | 10/54 |  |
| Total |  | Resident<br>Short-range nom.<br>Long-range nom. | 24/168 (0.14<sup>c</sup>)<br>19/179 (0.11)<br>30/250 (0.12) | 10/54 (0.19)<br>18/99 (0.18)<br>5/62 (0.08) | 7/52 (0.13)<br>3/18 (0.17) |

<sup>a</sup> Nomads. <sup>b</sup> Positive/tested. <sup>c</sup> Proportion of positive results

We computed 95% confidence intervals for prevalences and odds-ratios, indicated using squared brackets. All the statistical analyses were conducted using R 3.6.1 [48].

### Epidemiological model

#### Model design and parameterization

The epidemiological system was considered at the pond scale. Considering that nomadic herders always settle close to ponds and that the flight capacities of both *Culex* and *Aedes* are highly variable, ranging from 50m to 50km depending on species, climatic and environmental conditions, host availability and experimental protocols [49], we assumed that mosquitoes emerge from ponds to seek hosts in a 2.5 km radius, corresponding to the putative area of influence of the pond in this region. We considered two closed vector populations (*i.e.* without dispersion from or to the epidemiological system): a *Cx‥ poicilipes* and an *Ae. vexans* population. Four host populations were included: cattle and small ruminants from sedentary and nomadic herds. Resident ruminant populations were assumed closed for renewal (no purchase of live animals, culled animals being replaced by animals born in the epidemiological system). However, nomadic ruminant populations were assumed to have a permanent renewal due to arrival of some breeders and the departure of others. Each vector population was represented by a compartmental S-L-I model (S: susceptible vectors, L: infected but non-infectious vectors -during the extrinsic incubation period, I: infectious vectors that may transmit RVFV to the susceptible hosts upon which they feed), and by an additional state variable (G) representing the proportion of infected eggs. Host populations were represented by a S-I-R model (S: susceptible animals, I: viraemic animals, which may transmit RVFV to vectors feeding upon them, R: immune animals), further stratified by age (with yearly age classes) and by sex/physiological status (males, empty females and gestating females).

Model dynamics represented three distinct processes (Fig. 2): the population dynamics (birth and death of vectors and hosts), the infection dynamics (RVFV transmission) and the population renewal (arrival and departure of nomadic breeders). The population dynamics of vectors was not explicitly represented in the model. Instead, we used the outputs (computed with a daily time step) of the entomological model (EM) elaborated by Tran et al., and parameterized using mosquito trapping data collected in the same study area and period (Fig. 2) [2]. This model reproduced the demographic dynamics of *Ae. vexans* and *Cx. poicilipes* mosquitoes around a temporary pond, based on the evolution of pond surface, rainfall, temperature and humidity.

**Figure 2.**
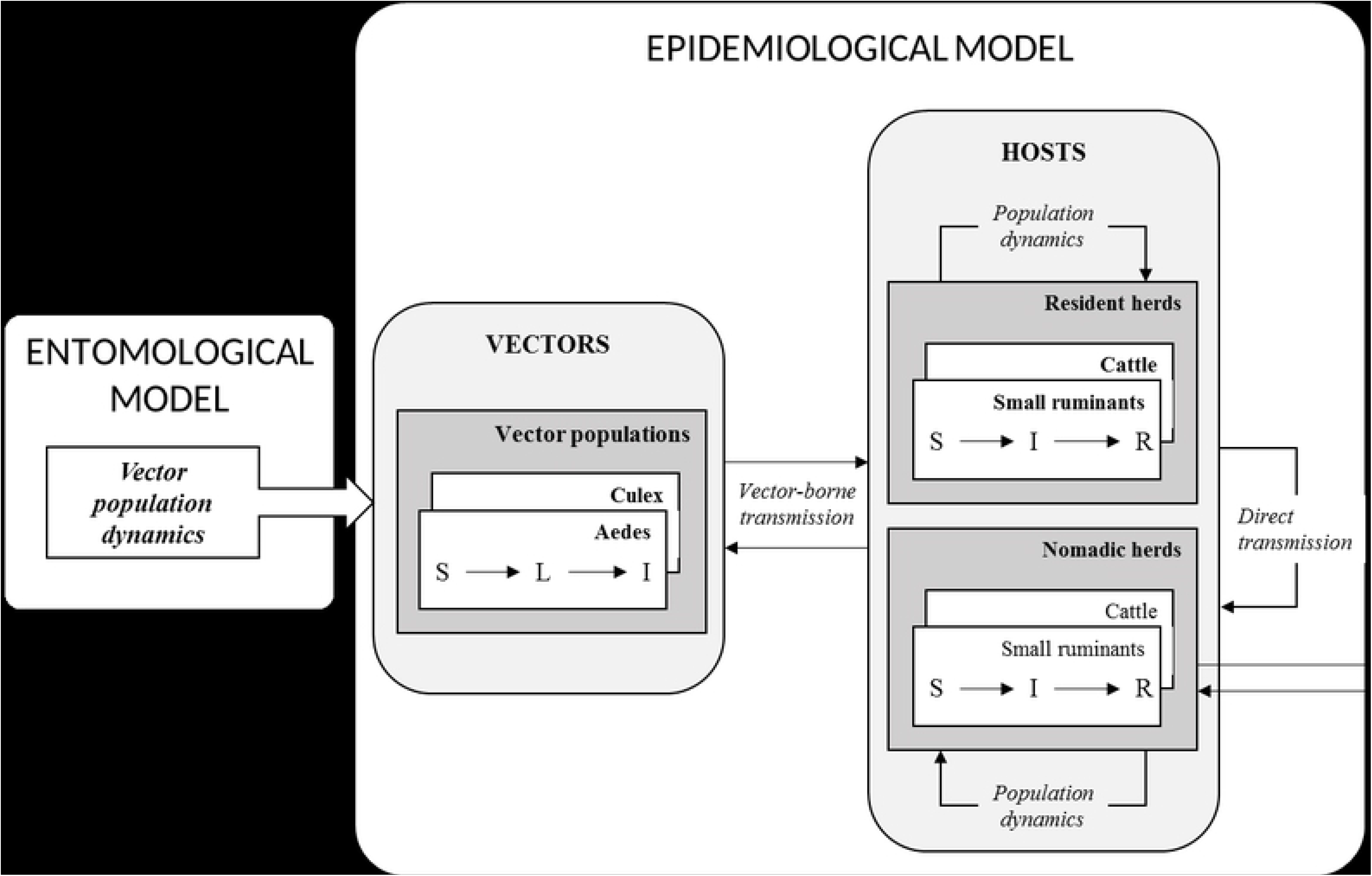
Schematic representation of the epidemiological model.

Two between-host transmission routes were considered: vector-borne, and direct, considering that when viraemic hosts abort (or calve/lamb), the abortion (or calving/lambing) products are infectious, and susceptible animals are exposed to these highly infectious materials (Fig. 2). Vector-borne transmission was parameterized by a scaling factor for vector population sizes (*ψ*), and direct transmission by a transmission parameter (*β*). In vectors, RVFV transmission from an infected female to its eggs was parameterized by a VT probability (*ω*).

Since the host and vector population sizes remained constantly high in the modelled system throughout the year (>500 individuals) and because the RVFV circulation level was also expected to be high due to the documented recurrence of RVFV transmission in the area, we chose to implement the model in a deterministic framework, with a discrete daily time step.

A full description of (i) the epidemiological model, (ii) its coupling with the entomological model, and (iii) its parameterization, are provided in supplement S2.

#### RVFV introduction scenarios and parameter estimation

The model was simulated during an 8 years duration corresponding to the maximal lifespan of hosts. Entomological inputs (vector population dynamics) were obtained from the EM for the 2008-2015 period, using daily rainfall, temperature and humidity data collected from the study area during this period. Two RVFV introduction scenarios were considered: a single introduction due to the arrival of a given number *n_intro_* of viraemic nomadic animals during the 2008 rainy season (scenario A), or successive introductions due to the arrival of *n_intro_* viraemic animals during each rainy season from 2008 to 2015 (scenario B). In both cases, the viraemic animals were proportionally allocated to cattle and small ruminants’ nomadic populations according to the species-specific numbers of susceptible individuals. For both scenarios, the risk of introduction and a consecutive initiation of transmission cycle was assumed to vary throughout a given rainy season, as (i) the geographic origin of nomadic breeders is known to vary throughout the rainy season [1], (ii) the succession in time of *Ae. vexans* (first part of the rainy season) and *Cx. poicilipes* populations (second part of the rainy season) [31] may also imply differences in RVFV circulation level, and (iii) a progressive amplification of RVFV circulation in the whole Ferlo area may induce an increased risk of introduction in the late rainy season. Therefore, the total number of viraemic animals entering the area during a given rainy season (*n_intro_*) was divided into (i)a proportion *p_early_* arriving during the first half of the rainy season, and (ii) 1 − *p_early_* arriving during its second half. Because of the daily time step, the number of viraemic ruminants entering the system at a given day was expected to be low (at least when the value of *n_intro_* is lower or comparable to the duration, in days, of the rainy period). To allow a realistic representation of such low numbers of viraemic animals entering the system, the day of arrival of each of the *n_intro_* ruminants was randomly chosen at the beginning of each simulation. This method introduced a stochastic component to the model, justifying the use of a Bayesian framework for parameter estimation, instead of likelihood maximization methods more commonly used for deterministic models. We used an Adaptive Population Monte-Carlo Approximate Bayesian Computation method (ABC-APMC) [50], with non-informative priors (assumed to be independent of one another) for each of the 5 estimated parameters:

- the scaling factor for vector population sizes (*ψ*), prior: U(0-10)
- the direct transmission parameter to hosts (*β*), prior: U(0-10)
- the probability that an infected *Aedes* female mosquito transmits the virus to its eggs (*ω_aedes_*, this probability being considered null for *Culex* females), prior: U(0-0.10) (this proportion was previously estimated at 0.007 [51])
- the total number of viraemic nomadic ruminants introduced during a rainy season (*n_intro_*), prior: U(0-1500)
- the proportion of viraemic nomadic ruminants arriving during the 1st half of the rainy season (*p*_*early*_), prior: U(0-1).

The summary statistics consisted of the observed numbers of seropositive resident small ruminants per age class (6 classes) found when simulating the serosurvey described above (i.e. during the 2016 dry season, with the age-specific numbers of tested animals given in Table 4). The recommended settings were used for the ABC-APMC algorithm: 0.5 for the quantile of the distribution of distances to observed data used to define tolerance thresholds; 0.05 for the stopping criterion based on the renewal proportion of particles in the sequential Monte-Carlo procedure; and a final size of 10,000 particles to build posterior probabilities [50].

The scenarios A and B were compared using a model selection procedure based on random forest classification methods, specifically designed to allow model comparison in an approximate Bayesian computation framework [52]. Using the same method, the need for VT and/or direct transmission between hosts was tested. Five models were compared (Table 3) and the best model was used for parameter estimation. When two models were equivalent, the most parsimonious was selected.

**Table 3.** Model selection: definition of the compared models.

| Model | Introduction scenario | Vertical transmission parameter in *Aedes* $\omega_{aedes}$ | Direct transmission parameter between hosts $\beta$ |
| --- | --- | --- | --- |
| M0 | A (2008 only) | Estimated | Estimated |
| M1 | B (2008-2015) | Estimated | Estimated |
| M2 | Best scenario* | 0 | 0 |
| M3 | Best scenario* | Estimated | 0 |
| M4 | Best scenario* | 0 | Estimated |

\* According to M0-M1 comparison

## Results

### Surveys in sedentary herds and in nomads

One hundred-sixty eight resident sheep and fifty four goats were sampled (Table 2 and 4). The overall IgG seroprevalence rate was 15.3%. [10.8-20.7], with 14.3 % [7.4-25.1] in Kodediaré, 10% [4.4- 20.1] in Nakara, and 20 % [12.9- 31.4] in Soringo. The IgG seroprevalence rate in goats was 18.5% [9.2- 31.4] and 14.3% [9.4-20.5] in sheep. We did not register any IgM positive sera. The seroprevalence rate increased significantly with age, determined based on the dentition of the animals (Table 4) (test for trend in proportions: p<0.005).

**Table 4.** Results of an age-structured serosurvey conducted in small ruminants from 6 resident herds and 22 nomadic herds of Younoufere area, Ferlo, Senegal, August 2015-May 2016.

| Age | Dentition | Positive/Tested<br>(proportion) |  |  |
| --- | --- | --- | --- | --- |
|  |  | Resident herds | Short-range<br>nomads | Long-range<br>nomads |
| 0-1 year | Milk teeth | 1/35 (0.03) | 2/61 (0.03) | 6/91 (0.07) |
| 1-2 years | 2 teeth | 2/36 (0.06) | 2/39 (0.05) | 3/56 (0.05) |
| 2-3 years | 4 teeth | 8/35 (0.23) | 13/57 (0.23) | 0/30 |
| 3-4 years | 6 teeth | 4/35 (0.11) | 9/43 (0.21) | 8/36 (0.22) |
| 4-5 years | 8 teeth | 10/39 (0.26) | 7/46 (0.15) | 9/42 (0.21) |
| >5 years | >8 teeth | 9/42 (0.18) | 4/32 (0.12) | 9/57 (0.16) |

An average number of 36 nomadic breeders per session were interviewed during the 3 surveys of the rainy season, and 12 breeders during the 2 surveys of the dry season (Table 1). Based on their declarations, nomadic herders would stay on average 16.3 days in Younoufere during the rainy season (range: 2-122 days), and 9.8 days during the dry season (range: 1-21 days). Herds were essentially composed of sheep with a variable number of cattle goats and sometimes donkeys. Four herds were dominated by cattle and two by goats. The distribution of the size of the 22 sampled herds as well as their compositions are provided in supplement S1.

A total number of 660 serological test results were available for the statistical analysis in nomadic animals (Table 2) among which 590 were from small ruminants and 70 from cattle (Table 4). The overall estimated IgG seroprevalence rate was 12.4% [10.0-15.2], specifically 14.3% in cattle (n=70; [7.4- 25.1]), 11.4% (n=429; [8.6-14.9]) in sheep, and 14.3% in goats (n=161; [9.5-20.9]). A higher seropositivity rate was detected in samples collected during the 2^nd^ half of the rainy season and the early dry season (Dec-Jan) than in samples collected during the 1^st^ half of the rainy season (Aug-Sept), with respectively 18.5% [14.1-23.8] and 8.2% [5.7-11.5] of sera detected IgG positive. The observed IgM seroprevalence was 0.5% [0.4-2.6] for herds sampled during the first half of the rainy season against 4% [2.1-6.9] for herd sampled afterwards.

Animal’s age was significantly associated with the seropositivity risk (OR: 1.3 [1.1-1.5] for an increase of 1 year of age). Small ruminants from nomadic herds (both long- and short-range) were more frequently seropositive when sampled during the 2^nd^ half of the rainy season (and the early dry season) than animals from long-range nomadic herds sampled during the 1^st^ half of the rainy season (the reference class). It was also the case for resident herds sampled during the 2016 dry season (Table 5).

**Table 5.** Logistic mixed model of the association between seropositivity with individual-level and group-level covariates in small ruminants from resident and nomadic herds of Younoufere area, Ferlo, Senegal, August 2015-May 2016.

| Variable | Value | p-value | Odds-ratio |
| --- | --- | --- | --- |
| Age | Quantitative (years) | <0.0001 | 1.3<sup>a</sup> [1.1-1.5] |
| Sex | Female | Reference |  |
|  | Male | 0.14 | NS |
| Species | Sheep | Reference |  |
|  | Goat | 0.25 | NS |
| Type of herd – sampling date | Long-range nom.<sup>b</sup> – 1<sup>st</sup> half of rainy season<sup>c</sup> | Reference |  |
|  | Short-range nom. – 1<sup>st</sup> half of rainy season<sup>c</sup> | 0.09 | NS |
|  | Long-range nom. – 2<sup>nd</sup> half of rainy season<sup>c</sup> | 0.0005 | 7.9 [2.3-28.3] |
|  | Short-range nom. – 2<sup>nd</sup> half of rainy season<sup>c</sup> | 0.009 | 4.5 [1.4-16.2] |
|  | Resident – Dry season<sup>d</sup> | 0.02 | 3.9 [1.2-14.7] |

<sup>a</sup> computed for an increase of 1 year of age, <sup>b</sup> Nomads, <sup>c</sup> 2015, <sup>d</sup> 2016

### Epidemiological model

Based on the posterior probability (>0.999), model M1 (yearly introductions of viraemic animals, scenario B) was first selected as compared to M0 (a single year with introduction of viraemic animals, in 2008) (Table 3). The comparison of model M1 (including both VT in *Aedes* and direct transmission) with model M2 (including none of these mechanisms) led to select model M1 (posterior probability 0.996). Then model M1 was selected against M3 (same as M1, without direct transmission; posterior probability 0.994). Lastly, M1 and M4 appeared equivalent (posterior probability: 0.32): we thus selected the most parsimonious model M4, which included direct transmission but no VT in *Aedes*, and repeated introductions of RVFV by nomadic animals, during each rainy season, from 2008 to 2015. Model fit obtained after the parameter estimation of model M4 appeared satisfactory (Fig. 3): the predicted age-specific distributions of IgG seroprevalence rate matched the results of the serosurvey in sedentary small ruminants, and the 95% confidence intervals of age-specific predicted distributions of seroprevalence rates always included the observed value (Fig. 3).

**Figure 3:**
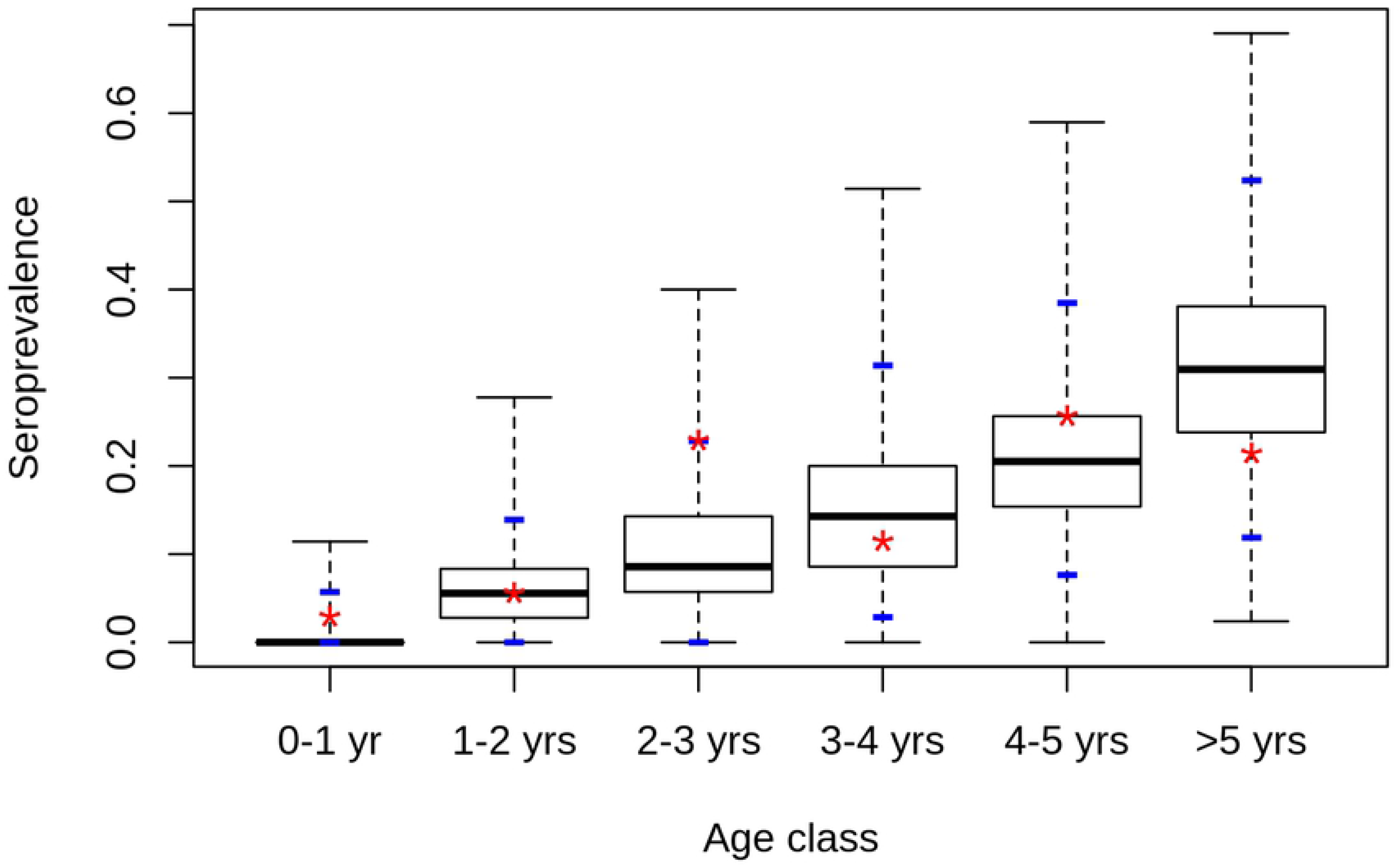
Predicted distribution of seroprevalence rate by age class in sedentary small ruminants. Model is model M4 which combines direct transmission between hosts but no VT in Aedes, and repeated introductions of RVFV by viraemic nomadic animals, each rainy season, from 2008 to 2015. Stars: observed values, blue dashes: 2.5% and 97.5% percentiles of predicted distributions

The posterior distributions of estimated parameters are provided in Fig. 4. The posterior distribution of the scaling factor for vector population sizes (*ψ*) had a median value of 0.09 (95% credibility intervals [CI]: 0.005-0.32). The median value of the direct transmission parameter (*β*), was 4.8 (95% CI: 1.8-9.3). This estimation is close to the estimate previously obtained in Madagascar highlands, by Nicolas et al. (3.2 [3.0-3.4]). The posterior median value of the total number of viraemic nomadic animals introduced during a rainy season (*n_intro_*) was 617 (95% CI: 211-1,336). The posterior median of the proportion of viraemic animals arriving during the 1^st^ half of the rainy season (*p_early_*) was 0.31 (95% CI: 0.02-0.88):according to the model, viraemic animals were more likely introduced during the late rainy season than early, which is consistent with a progressive amplification of RVFV in the area.

**Figure 4.**
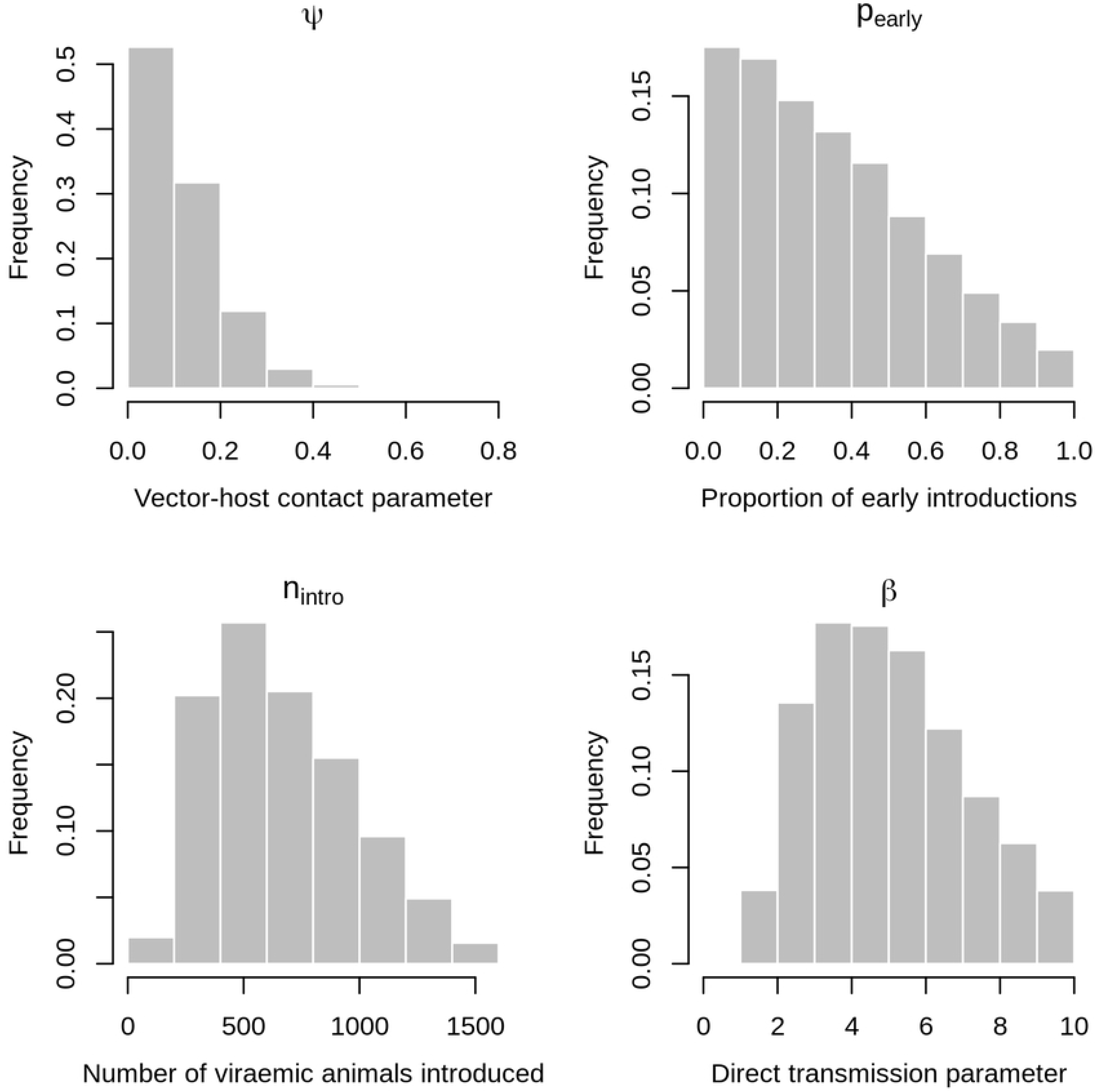
Posterior distributions of estimated parameters. Model is M4 which combines direct transmission between hosts but no VT in *Aedes*, and repeated introductions of RVFV by viraemic nomadic animals, each rainy season, from 2008 to 2015. *ψ*: scaling factor for vector population sizes, *β*: direct transmission parameter between hosts, *n_intro_*: total number of viraemic nomadic animals introduced during a rainy season, *p_intro_*: proportion of those animals arriving during the 1^st^ half of the rainy season, *p_early_*: proportion of viraemic nomadic ruminants arriving during the 1st half of the rainy season

More precisely, combining the two preceding posterior distributions resulted in a median number of 172 (95% CI: 8-742) and 390 (95% CI: 48-1,097) viraemic animals introduced during the 1^st^ and 2^nd^ halves of the rainy season, respectively. Moreover, a total number of 22,686 nomadic animals (3,796 cattle and 18,890 small ruminants) entered the study area during the 2015 rainy season, of which 50% (11,343 animals) arrived in the first half of the rainy season and 50% during the second half of the rainy season. The predicted proportions of viraemic animals among nomadic ruminants entering the study area in 2015 had then a median value of 1.5% [0.1-6.5] for the 1^st^ half of the rainy season and 3.4% [0.4-9.7] for the 2^nd^ half. These predicted proportions are consistent with observed IgM seroprevalence rates of 0.5% and 4.0% observed in nomadic animals entering the study area during the 1^st^ and 2^nd^ halves of the 2015 rainy season, respectively.

## Discussion

RVF has been repeatedly reported in northern Senegal since 1987, when the first large outbreak of west Africa started in southern Mauritania [45, 53, 54]. Since then, 5 outbreaks were reported by the national RVF surveillance network in 2003, and a survey showed an active viral circulation with multiple clinical cases in small ruminants in the Ferlo area [25]. A longitudinal serosurvey carried in the same area during the 2004 rainy season confirmed a recurrent RVFV transmission [55]. More recently, RVFV was isolated from *Ae ochraceus* in the same area thanks to a mosquito-based surveillance implemented since 1990 [46]. The last evidence of RVFV circulation occurred during the 2013-2014 outbreak with the detection of one human case in Linguere town located in northern Senegal [56]. Our serological results confirm a long lasting RVF endemicity in resident herds. As expected, we did not detect IgM because residents animals were sampled during the dry season. This survey also provides the first estimation of RVF IgG seroprevalence in nomadic herds in West Africa, which was close to what was estimated in resident herds. In small ruminants, whatever the type of herd (resident, short-range nomad and long-range nomad), seroprevalence data increases with age. Despite the small sample size, these results suggest that RVF force of infection is equally high in both nomadic and resident herds because of an endemic circulation of RVFV in the whole Ferlo region and a consecutive regular exposition of herds to the virus. In addition, the multivariate analysis (logistic mixed model), showed that, besides the increase of seropositivity risk with age, the seropositivity risk increased between the 1^st^ and 2^nd^ half of the rainy season. This suggests an amplification of the transmission cycle during the rainy season with a peak of circulation at the end of that season. This latter assumption is reinforced by an observed IgM seroprevalence of 0.5% for nomadic herds sampled during the 1^st^ half of the rainy season, against 4% (CI 2.1-6.9) for herds sampled on late rainy season, and the isolation of a single RVFV strain in the same area in September 2012 (2^nd^ half of the rainy season [46]). The observed IgM rates show that RVFV has been circulating within the Ferlo area between August 2015 and December 2016, and, similarly to what has already been observed in many endemic countries [57–59], without any reported clinical case, either in ruminants nor humans. Lastly the observed IgM rate is in line with previous results reported in other endemic areas, i.e. 0.5% in Tanzania, 0–4% in Mayotte and 1.3% recently in South Africa [57, 60, 61].

Given the absence of mosquito vectors during the dry season, the way the virus is maintained in the area remains unknown. Here, we assumed that RVF transmission recurrence may be explained by a source-sink dynamic, at least at the Ferlo scale, nomadic herds constantly moving from ponds to ponds according to the availability of water, thus introducing occasionally or seasonally the virus in epidemiological systems where the absence of mosquitoes during the dry season prevents a yearlong virus transmission. To investigate this hypothesis, we modelled RVF transmission at the pond scale, during an 8 year time period (corresponding to the average life span of ruminants), incorporating both *Ae. vexans* and *Cx. poicilipes* population dynamics, resident and nomadic ruminant population dynamics and movements. Five models were compared that combined a single or yearly introductions of RVFV with the presence or absence of VT and of direct transmission. The selected model incorporated yearly introductions of RVFV from 2008 to 2015 (the study period) by nomadic herds, with a larger proportion of viraemic animals arriving during the 2^nd^ half of the rainy season. This result is consistent with the assumption of an amplification of transmission during the rainy season. The predicted proportion of viremic animals introduced during the 2015 rainy season is also in line with the IgM rate observed in nomadic herds, sampled during either the first (0.5%) or the second half of the rainy season (4%). In addition, since IgM antibodies do not persist beyond the 50^th^ day after infection in the majority of cases [62], and since the majority of IgM positive animals were sampled in December-January, the infection of these animals likely occurred in November-December, thus confirming the amplification of RVFV circulation at the end of the rainy season.

Implication of vertical transmission (VT) in RVFV persistence in a given area during inter-epizootic periods remains widespread cited. In our study, the selected model did not incorporate VT mechanisms, suggesting that (i) VT is not necessary to account for RVFV recurrence in the Ferlo area, and (ii) nomadic herd movement migration are sufficient to account for a re-initiation of the transmission cycle every year. As a matter of fact, the implication of VT in the persistence of the virus in the field during unfavourable seasons, only relies on few experimental or empirical observations. First, RVFV was detected in females and males *Ae. mcintoshi*, reared in laboratory from field collected eggs in Kenya [37]. However, and to our knowledge, these findings have never been reported again, even during epizootic periods, and whatever the geographic area. RVFV genome was further detected in adult male *Ae. vexans* and *Cx. quinquefasciatus* in Sudan in 2009-2010. However, the presence of infectious particles has never been confirmed [63]. The last argument relies on the fact that several outbreaks occurred at the same time in regions separated by hundreds of kilometres, but concurrently exposed to heavy floodings that may have triggered a massive *Aedes* population hatching, some of these *Aedes* being possibly infectious [36]. However, such heavy flooding also attract nomadic herds and provoke mosquito population explosions, both favourable to RVFV spread, transmission and animal or human RVF case occurrence after several years of a low-level virus circulation.

The selected model included a direct transmission mechanism between hosts to take into account the excretion of RVFV in abortion products (or lambing/calving products) of viraemic females, that may be a source of infection for other ruminants (as they are for humans). The estimate of the direct transmission parameter was close to the value obtained by Nicolas et al [35] in a very different ecosystem (Madagascar highlands). Although limited to specific conditions (abortion or calving/lambing of viraemic animals), this direct transmission mechanism may thus play a significant role in the local RVFV circulation, while being independent of the eco-climatic context.

RVFV was recently isolated from *Ae. ochraceus* [46], and several entomological surveys showed the presence of many potential RVFV vectors such as *Mansonia uniformis* or *Cx. tritaenhyorynchus* in the Ferlo. However *Ae. vexans* and *Cx. poicilipes* are considered the main vectors of RVF in this region [26, 30]. Their respective population dynamics was first described by Mondet et al in 2002 [31], with an early rainy season characterized by an abundant population of *Ae. vexans*, whereas the second part of the rainy season was favorable to *Culex* population. These observations have been recently confirmed [38], with *Ae. vexans arabiensis* representing 94.98% of mosquitoes captured, and *Cx. poicilipes* 0.24% of the mosquito trapped. Despite this low proportion of *Cx. poicilipes*, we chose to incorporate both vector species in our model, to account for seroconversions occurring late in the rainy season, when *Ae. vexans* is absent.

The number and size of nomadic (and sedentary) herds were assumed constant during the study period, although the duration of the rainy season varies from year to year, which may increase or decrease the number and size of nomadic herds. We made the hypothesis that the duration of the 2015 rainy season was not very different from the average, as confirmed by the estimated dates for the beginning and end of the 2008-2014 rainy seasons (Table S4), and that the ruminant population dynamics parameters observed in 2015 were close to the usual conditions prevailing in the study area.

Outbreaks of vector-borne diseases such as RVF are known to be sensitive to both host movements and landscape characteristics, with factors such as host densities and movement patterns contributing to disease maintenance [35, 64, 65]. RVF is endemic in the Ferlo area. In line with Favier et al [66] who suggested that (i) vertical transmission or wild reservoirs are not necessary to explain RVFV endemicity in the Ferlo, (ii) herd movements in some specific environment can allow endemicity at a regional scale while circulation is epidemic at a local scale, we quantitatively demonstrate in this work that although the existence of a vertical transmission mechanism in *Aedes* cannot be ruled out, nomadic movements are sufficient to account for this endemic circulation in the Ferlo area.

A next outbreak will inevitably occur in the Ferlo area, and the increased national and transboundary nomadic and commercial movements of ruminant herds may allow the virus spreading over large distances, threatening disease-free areas. It is thus urgent to improve the surveillance capacity as well as our knowledge on nomadic herds, their movements and the determinants of these movements. Since they should be targeted in any vaccination campaign, it is finally necessary to learn more about the perception of RVF by nomadic breeders, and the social acceptability of prevention, surveillance, and control measures, such as cattle, small ruminant and dromedary vaccination to protect people, or animal movement restrictions to avoid RVFV spread through hubs in the livestock-trade networks

## Acknowledgments

This study was supported by Vmerge project (Emerging viral vector-borne diseases; https://www.vmerge.eu/) and by the Ile-de-France Region as part of the DIM-1Health project. We gratefully thank Younoufere authorities and breeders

## Supporting information

S1. Characteristics of the sampled nomadic herds

S2. Epidemiological model

S3. Checklist: STROBE Checklist

